# High-quality metagenome assembly from nanopore reads with nanoMDBG

**DOI:** 10.1101/2025.04.22.649928

**Authors:** Gaëtan Benoit, Robert James, Sébastien Raguideau, Georgina Alabone, Tim Goodall, Rayan Chikhi, Christopher Quince

**Author notes:** These authors jointly supervised this work (Rayan Chikhi, Christopher Quince).

## Abstract

Third-generation long-read sequencing technologies, have been shown to significantly enhance the quality of metagenome assemblies. The results obtained using the highly accurate reads generated by PacBio HiFi have been particularly notable yielding hundreds of circularized, complete genomes as metagenome-assembled genomes (MAGs) without manual intervention. Oxford Nanopore Technologies (ONT) has recently improved the accuracy of its sequencing reads, achieving a per-base error rate of approximately 1-2%. Given the high-throughput, convenience and low-cost of ONT sequencing this could accelerate the uptake of long read metagenomics. However, current metagenome assemblers are optimized for PacBio HiFi data and underperform on the latest ONT data and do not scale to the large data sets that it enables.

We present nanoMDBG, an evolution of the metaMDBG HiFi assembler, designed to support newer ONT sequencing data through a novel pre-processing step that performs fast and accurate error correction in minimizer-space. Across a range of ONT datasets, including a large 400 Gbp soil sample sequenced specifically for this study, nanoMDBG reconstructs up to twice as many high-quality MAGs as the next best ONT assembler, metaFlye, while requiring a third of the CPU time and memory. As a result of these advances, we show that the latest ONT technology can now produce results comparable to those obtained using PacBio HiFi sequencing at the same sequencing depth.

## Introduction

Metagenomics, the sequencing of genetic material recovered directly from environmental samples, enables the exploration of complex microbial communities [27, 23]. The initial step in metagenomic analysis is the assembly of sequencing reads into longer contiguous fragments or contigs. Metagenome assembly from short-read sequencing poses significant challenges, often yielding highly fragmented assemblies, particularly in diverse communities. Although binning methods can cluster short-read contigs into metagenome-assembled genomes (MAGs) [1], MAGs constructed from short reads frequently suffer from incompleteness, contamination and high fragmentation.

Long-read sequencing technologies, Oxford Nanopore Technologies (ONT) and PacBio HiFi, have significantly enhanced the quality of metagenomic assemblies. PacBio HiFi reads, characterized by their high accuracy, have made it possible to resolve hundreds of circular genomes directly from complex metagenomic samples, offering unprecedented insights into microbial communities [4, 11, 3]. While applications of earlier ONT reads to metagenomics were limited by high error rates [18], recent advances in ONT chemistry and base-calling algorithms now enable reads with error rates around 1% [32, 31], opening new avenues for methods development.

MetaFlye was the first pioneering assembler specifically developed for long-read metagenomic assembly [13]. This was designed to tackle the assembly both of noisy ONT data and HiFi. Hifiasm-meta in contrast, specifically harnesses the high accuracy of HiFi reads to reconstruct complete, circularized genomes and outperforms metaFlye in this regard. More recently we introduced metaMDBG, also optimized for assembling HiFi metagenome data, which reduces memory usage by an order of magnitude compared to hifiasm-meta and surpasses all current tools in both the production of near-complete MAGs and runtime efficiency. However hifiasm-meta and metaMDBG both underperform on ONT data, even with the latest base-calling accuracy improvements. As a result, metaFlye remains the only viable option for assembling ONT datasets. This gap motivated us to develop an evolution of metaMDBG specifically designed to address the error handling challenges in ONT data.

At the core of our algorithms is a variant of the de-Brujin graph (DBG) tailored for long reads, the minimizerspace de-Brujin graph (MDBG), first introduced in rust-mdbg then refined in metaMDBG [10, 3]. This approach considers only a small subset (around 0.5%) of k-mers from each read, selected uniformly using a minimizer-based sampling technique. The chosen k-mers are then linked into chains of size *k*^*′*^ and further extended using a de Bruijn graph-like framework. In order to exploit the full potential of high-accuracy long reads, metaMDBG creates long and specific chains (k’ *≈* 100). However in the presence of sequencing errors, such long chains inevitably end up containing erroneous k-mers which break contiguity.

A first attempt to deal with sequencing errors in minimizer-space was introduced in rust-mdbg [10], through read correction inspired by multiple sequence alignment. For each read represented by its minimizers, similar reads with a high fraction of shared minimizers are recruited then aligned to generate a consensus sequence in minimizer-space using partial order alignment (POA). This method has notable limitations. Firstly, the read recruitment implemented using a resource-intensive disk-based read bucketing step degrades performance. Secondly, a low similarity threshold for read recruitment is overly permissive, especially in the metagenomics context, leading to reads from different species being potentially aligned together which in turn yields erroneously corrected reads. While a higher similarity threshold could mitigate this risk, it would also exclude genuine alignments with long flanking regions. Also, the low density of minimizers typically utilized by rust-mdbg and metaMDBG precludes accurate estimation of divergence between reads and leads to subpar recruitment.

We introduce nanoMDBG, which revisits the concept of read correction in minimizer-space to address errors in ONT data. At the heart of nanoMDBG is an innovative use of fast seed-and-chaining techniques for read correction. It enables the rapid alignment and pileup of minimizers from similar reads to efficiently build consensuses. We further optimize the resource-intensive all-vs-all read mapping phase, a common bottleneck in read correction, by leveraging variable minimizer densities. Specifically, a low-density setting allows for quick recruitment of the most similar reads needed for effective correction, while a high-density setting ensures precise estimation of sequence divergence among the recruited reads.

This improved error correction allows nanoMDBG to handle large ONT datasets efficiently and accurately. For instance, on the largest tested dataset, a 400 Gbp Soil sample, the correction only took 16 hours using 32 cores (8% of overall assembly time) and 74 GB of memory (a fraction of the subsequent assembly requirements). On this dataset, nanoMDBG significantly outperformed state-of-the-art assemblers, recovering 244 more near-complete MAGs than metaMDBG and 180 more than metaFlye. When using PacBio HiFi data from the same samples, we demonstrate that nanoMDBG can now deliver ONT-derived results comparable in quality to those achieved with HiFi sequencing.

### Overview of minimizer-space assembly with nanoMDBG

We present nanoMDBG, a metagenome assembly method designed for Oxford Nanopore Technologies (ONT) long-read data. Given a set of input reads, nanoMDBG produces a FASTA file containing assembled contigs. An overview of the assembly workflow is illustrated in Figure 1. The universal minimizers, which are k-mers that map to an integer below a fixed threshold (see Methods), are first identified in each read. Each read is thus represented as an ordered list of the selected minimizers, denoted as an mRead. To improve accuracy, mReads are corrected using a multiple sequence alignment approach, employing a seed-and-extend strategy to align minimizers. For a given target mRead to correct, up to twenty of the most similar mReads are recruited, using a low-density density of minimizers to speed-up the alignment process (20% of the minimizers in each mRead by default). The recruited mReads are then re-aligned to the target mRead using the complete minimizer set, refining divergence estimation and reducing recruitment of reads from unrelated genomes. These alignments are then used to pileup minimizers using a minimizer-space variation graph, and a consensus sequence is extracted from the most supported path. The corrected mReads are then assembled into mContigs using the metaMDBG assembler. These mContigs are then converted back into base-space and polished.

**Figure 1.**
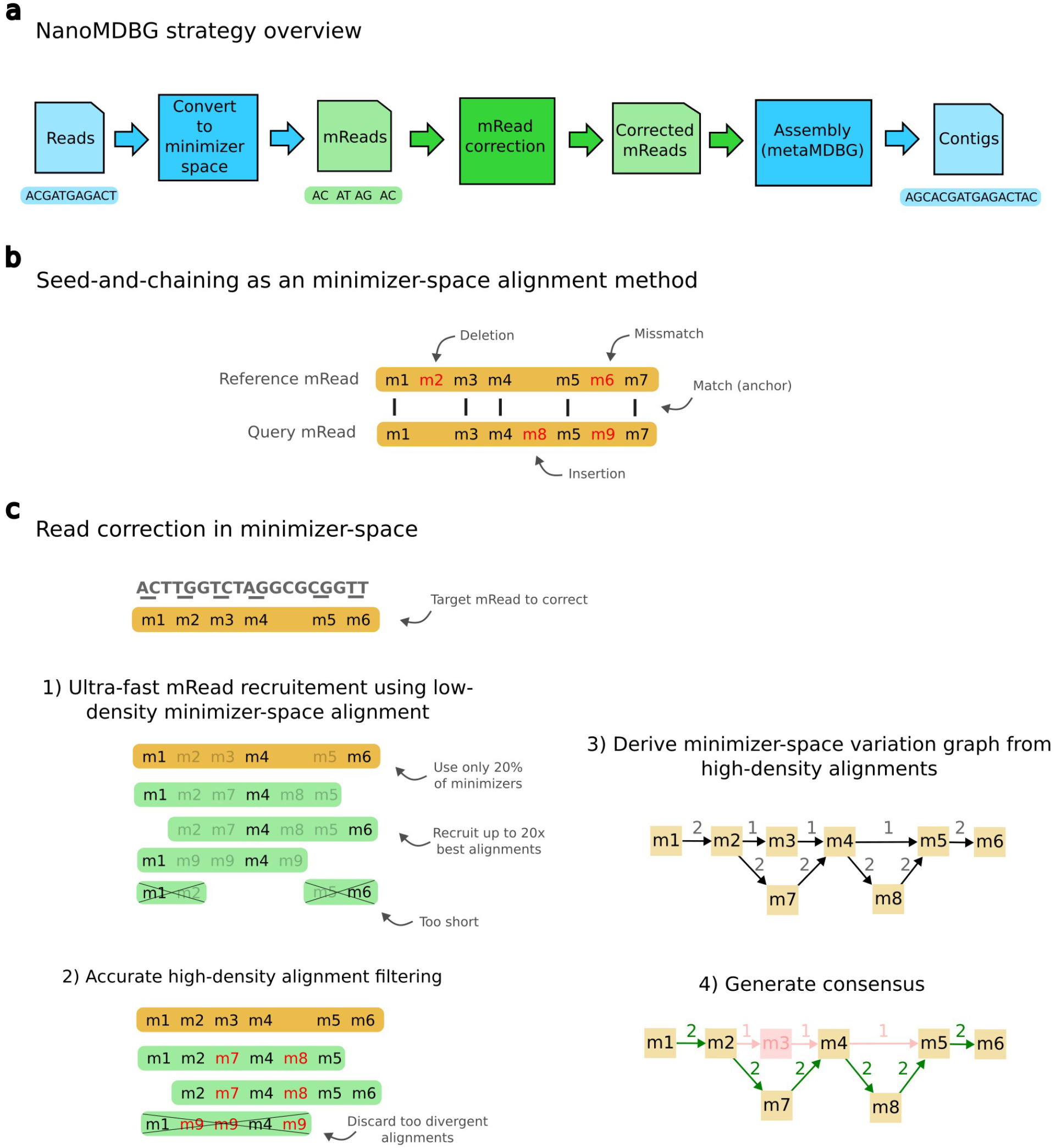
Overview of the algorithmic steps of nanoMDBG. **A**, Overview of nanoMDBG workflow. Steps in green correspond to our new contributions. **B**, Illustration of an alignment at the level of minimizers. **C**, Overview of the read correction method in minimizer-space. **1**, For a given target read to correct (in yellow), the most similar reads are recruited (up to 20x coverage) using seed-and-chaining alignment. We use only 20% of the read minimizers (by lowering the threshold for selecting univeral minimizers) to speed-up this costly recruitment process. **2**, Using the full read minimizers, we re-align the recruited reads using seed-and-chaining in order to accurately estimate their divergence with the target read. Reads with divergence over 4% are discarded. **3**, Using the accurate alignment computed in the previous step, we pileup the minimizers using a simple variation graph. The graph is initiated with the minimizer from the target read (path on top of the graph). Recruited reads are then added sequentially. A mismatch in a recruited read create a path parallel to a node from the target read (node m7), while an insertion create a path between two nodes from the target read (node m8) **4**, The most supported path (in green) is extracted from the graph and used as a consensus for the target read.

## Results

We first evaluate assembly results on ONT data only. We then compare assemblies obtained from ONT and HiFi data generated from the same samples.

### Evaluation of nanoMDBG for ONT metagenome assembly

#### Benchmarking setup

We compared nanoMDBG (v1.1) to two state-of-the-art assemblers: metaFlye (v2.9.3-b1797) and metaMDBG (v1.1) on one mock community and three real metagenomes (see Table 1). The commands that were used are provided in Supplementary Table S1 and all assembly results are summarized in Supplementary Table S3. A comparison to hifiasm-meta (v0.3-r063.2) is also given, although only on a subset of the data sets as explained below. The mock community, ‘Zymo’, contains 21 species for which reference genomes and abundances are known. The first real metagenome, ‘Human Gut’, is a 50 Gbp Human gut sample. The second metagenome, ‘Zymo Fecal Reference’ is a 200 Gbp reference standard constructed by pooling fecal samples from multiple donors [24]. The third data set, ‘Soil’, is a 400 Gbp soil sample taken from an agricultural field in Oxfordshire (UK) - see Methods. All data sets were sequenced with the latest R10.4.1 ONT flow cells and reads subsampled after filtering by selecting the first N reads necessary to achieve the specified data set sizes. This was done to allow comparison with equivalent sized HiFi PacBio data sets below. We note that the Human Gut and Soil datasets were generated for this study, while the Zymo Fecal Reference dataset is publicly available.

**Table 1.** Evaluated metagenome datasets. Two samples were previously publicly available (ONT Zymo Fecal Reference and HiFi Zymo Fecal Reference), and we newly sequenced five samples (ONT Zymo, ONT Human gut, ONT Soil, HiFi Human gut and HiFi Soil). To facilitate comparison across sequencing technologies, these datasets were subsampled by selecting the first N reads until the desired data volume was reached. Statistics were compute with command ‘seqkit stats –all’ [34].

| Sample | Accessions | #<br>bases<br>(Gb) | N50 read<br>length<br>(kb) | Average<br>quality<br>score | Sample description |
| --- | --- | --- | --- | --- | --- |
| Oxford Nanopore datasets |  |  |  |  |  |
| Zymo | - | 12 | 5.9 | 21.2 | Mock community Zymo-BIOMICS D6331 |
| Human gut | - | 50 | 7.8 | 19.6 | Human gut sample |
| Zymo Fecal Reference | - | 206 | 12.2 | 17.4 | Reference standard constructed by pooling fecal samples from multiple donors |
| Soil | - | 400 | 9.8 | 20.5 | Agricultural field in Oxfordshire (UK) |
| HiFi PacBio datasets |  |  |  |  |  |
| Human gut | - | 50 | 8.9 | 27 | Human gut sample |
| Zymo Fecal Reference | - | 255 | 8 | 28.8 | Reference standard constructed by pooling fecal samples from multiple donors [24] |
| Soil | - | 250 | 10 | 26.4 | Agricultural field in Oxfordshire (UK) |

#### Evaluation of bacterial genome reconstruction

We first evaluated the assemblers on the mock community Zymo, by aligning contigs to references and computing average nucleotide identity (see Methods). The results are summarized in Supplementary Table S2. Hifiasm-meta produced a highly fragmented assembly, likely because it is optimized for high-accuracy PacBio HiFi data; therefore, we excluded it from further comparisons in this section. NanoMDBG and metaFlye performed similarly, both in terms of the number of species recovered as circularized contigs and the ANI to reference sequences (*>*99.98% in most cases). MetaMDBG generally exhibited slightly higher fragmentation. The Zymo mock community contains 21 genomes, but five have very low coverage and five are strains of E. coli. In this case, nanoMDBG and metaFlye both obtained five circularized genomes and metaMDBG obtained four. No assembler could correctly resolve all of the E. coli strain diversity; however, nanoMDBG produced a consensus contig of 4,586,643 bp aligned to the reference strain B1109 with 99.79% ANI, covering 96.6% of the reference genome length. In contrast, metaFlye produced only fragmented genomes. Other abundant species were reconstructed in 1 to 6 linear contigs by nanoMDBG and metaFlye.

In the case of the real microbiome data sets data we used SemiBin2 [22] (v.2.1.0) to bin contigs obtained by each of the assemblers and evaluated the quality of the resulting bins with checkM2 [9] (v.1.0.1). We grouped bins into conventional categories based on their level of completeness and contamination reported by checkM2: near-complete MAGs (*>*90% completeness and *<*5% contamination), high-quality MAGs (*>*70% completeness and *<*10% contamination) and medium-quality (*>*50% completeness and *<*10% contamination). We also report the number of near-complete MAGs in a single contig (scMAGs) after binning.

For all three tested real communities, nanoMDBG reconstructed significantly more near-complete MAGs than the other assemblers (see Figure 2a). From the human gut sample, nanoMDBG reconstructed 80 near-complete MAGs (30 more than metaMDBG and 21 more than metaFlye), 235 from the Zymo Fecal Reference dataset (56 more than metaMDBG and 68 more than metaFlye) and 296 from the soil sample (244 more than metaMDBG and 180 more than metaFlye). The number of scMAGs generated by nanoMDBG is substantially improved compared to the other assemblers. NanoMDBG reconstructed 27 scMAGs from the human gut sample (16 more than metaFlye), 99 from the Zymo Fecal Reference dataset (55 more than metaFlye) and 158 from the Soil sample (146 more than metaFlye). On the Soil dataset, nanoMDBG reconstructed more complete genomes in a single-contig than the total number of near-complete MAGs found by metaFlye. As a further validation of the quality of the near-complete MAGs, we predicted the presence of rRNA and tRNA genes (see Methods). The near-complete MAGs generated by each assembler usually do contain the expected complement of RNA genes (91% for nanoMDBG and metaFlye, and 89% for metaMDBG). We also note that the majority of the near-complete MAGs produced by all assemblers contains fewer than 10 contigs (see Figure S1).

**Figure 2.**
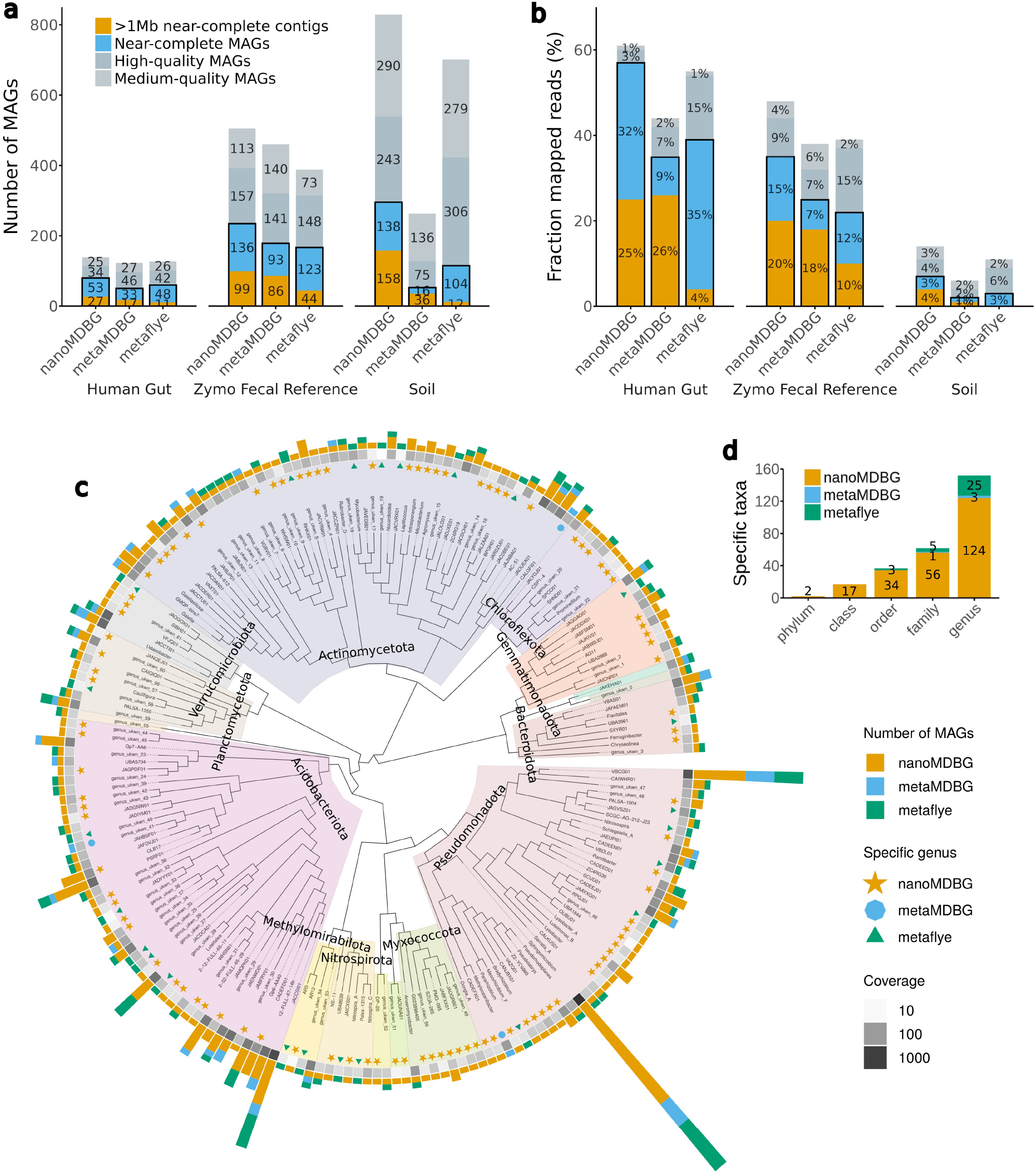
Assembly results on the three ONT metagenomics data sets. **a**, CheckM evaluation. A MAG is ‘near-complete’ if its completeness is *≥*90% and its contamination is *≤*5%, ‘high-quality’ if completeness *≥*70% and contamination *≤*10%, ‘medium quality’ if completeness *≥*50% and contamination 10%. **b**, The percentage of mapped ONT reads on MAGs. **c**, Phylogenetic tree of genera recovered from the ONT soil data set for all assemblers combined. For the near-complete bacterial MAGs, we generated a de novo phylogenetic tree based on GTDB-Tk marker genes, displayed at the genus level. The outer bar charts give the number of MAGs found in each genus. The colored symbols then denote genera recovered by only one of the assemblers. The grayscale heat map illustrates the aggregate abundance of dereplicated MAGs in a genus. **d**, Number of taxa at different levels that are unique to each assembler.

The improvement in assembly quality by nanoMDBG resulted in MAG collections that mapped a larger fraction of reads from each dataset (see Figure 2b) and hence, are more representative of the community. For the human gut dataset, nanoMDBG was the only assembler to map more than 50% of reads to near-complete MAGs. As the complexity of the datasets increased, the fraction of mapped reads decreased. On the Zymo Fecal Reference 35% of reads were mapped to near-complete MAGs but only 7% on the Soil near-complete MAGs. Not every read will be represented in the assembly, we should therefore compare these percentages to the fraction of reads that map onto the entire assembly, which gives their upper limits, these were Gut: 84%, Zymo Fecal Reference: 78% and Soil: 49%. Even accounting for the fact that not all genomes will be from prokaryotes this suggests that we are underperforming in the Soil data set.

To further investigate whether the low fraction of mapped reads in the Soil dataset was due to insufficient sequencing depth or assembly failure, we compared the number of recovered MAGs to the actual species diversity in the assembly. This was estimated by identifying single-copy core genes (SCGs) from contigs and clustering them at 97% identity (see Methods) (see Figure 3). Based on SCG analysis, we estimated that the human gut dataset contained 132 species, the Zymo Fecal Reference dataset 485 species, and the soil dataset 4,157 species. In the human gut dataset, 70% of predicted species had high coverage (*>*10x), and 85% of those were converted into MAGs. In the Zymo Fecal Reference dataset, over 90% of abundant species (*>*10x coverage) were also converted into MAGs. However, in the Soil dataset, only 36% of predicted species had coverage *>*10x, and just 39% of those abundant species were converted into MAGs. The low fraction of mapped reads in the Soil dataset can therefore be attributed to both the large species richness and the limited sequencing depth relative to that diversity, as well as challenges in assembling abundant genomes.

**Figure 3.**
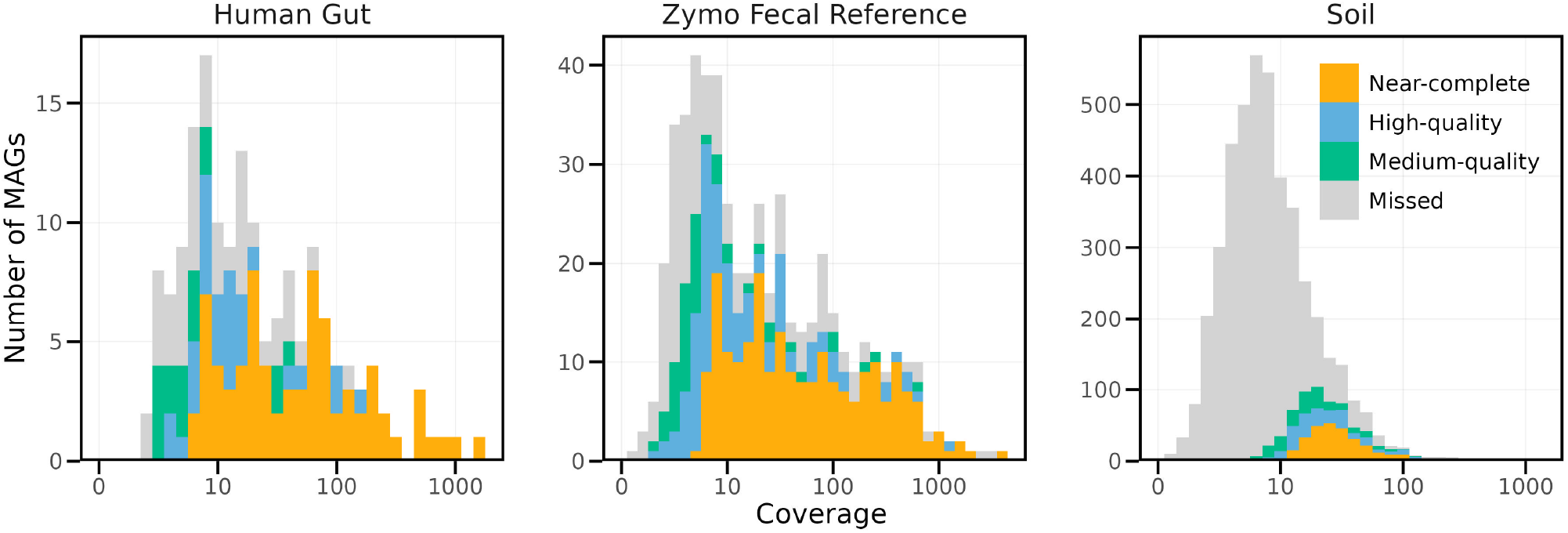
Rank abundance plot of species identified across the three ONT samples. Species counts were estimated based on the presence of the single-copy core gene COG0060 in nanoMDBG contigs, based on 97% identity clustering (see Methods). Colors indicate MAG quality, with “missed” representing species where the SCG was detected but not assigned to a MAG.

To summarize the microbial diversity from the soil sample, we constructed a phylogenetic tree with leaves as genera (see Methods) for all near-complete MAGs from all assemblers (see Figure 2c). We note that amongst the total 447 retrieved near-complete MAGs, only 5 could be assigned to a known species with a reference genome (ANI *>* 95%). The improved MAG recovery by nanoMDBG translates into a more representative picture of microbial diversity at all levels of evolutionary divergence. In total, we observed 98 genera that were recovered from the soil data sets by nanoMDBG but are missing from the near-complete MAG collections of the other programs. When the other assemblers did recover MAGs from the same genus, nanoMDBG usually found more MAGs. Finally, we can see large parts of the tree that are represented by only nanoMDBG MAGs; indeed, one phyla (38 families) were found only by nanoMDBG, compared to 5 families specific to metaFlye and one family specific to metaMDBG (see Figure 2d and Supplementary Table S6).

#### Evaluation of phages and plasmids reconstruction

We used geNomad [6] to identify circular contigs that were potential plasmid and phage genomes (see Supplementary Table S4). On the Gut and Zymo Fecal Reference samples, metaFlye reconstructed more viruses than nanoMDBG (3 and 58 more respectively), but nanoMDBG found 292 more phages in the soil data set. We used checkV [20] to assess the completeness of circular phages. We found that 60% of genomes recovered by nanoMDBG were judged high-quality against 57% for metaFlye. NanoMDBG found substantially more circular plasmids compared to metaFlye for all three metagenomes (14 more on the human gut sample, 37 more on the Zymo Fecal Reference and 7 more on the Soil sample). MetaMDBG usually obtained less circular viruses and plasmids than nanoMDBG and metaFlye.

#### Evaluation of NanoMDBG scalability

By performing the correction step in minimizer-space, our novel method, nanoMDBG, naturally extends metaMDBG to ONT data without significantly affecting overall performance. For example, on the 400 Gbp Soil sample, the correction step required only 16 hours using 32 cores (accounting for 8% of the total assembly time) and 74 GB of memory, which is a fraction of the memory required for subsequent assembly (see Supplementary Table S5). NanoMDBG completed the full soil dataset assembly in just 9 days, compared to 28 days for metaFlye. In terms of memory efficiency, nanoMDBG used 211 GB, while metaFlye required 753 GB, making the latter potentially inaccessible for many laboratories. Both assemblers efficiently handled the 50 Gb Human gut sample, with nanoMDBG completing the task in 5 hours and metaFlye in 8 hours. However, nanoMDBG used only 20 GB of memory compared to metaFlye’s 115 GB, enabling nanoMDBG to potentially assemble datasets of equivalent complexity on a laptop, which could be important given the portability of ONT sequencing devices.

#### Comparison with base-level read correction tools

We explored an alternative approach for processing ONT data by applying read correction tools prior to assembly. Specifically, we assessed two recent haplotype-aware correction tools: HERRO [35] and DeChat [15]. While DeChat is tailored for metagenomics, we note that HERRO was developed for genomic data, with the authors recommending its use on reads longer than 10 kbp, significantly higher than the typical N50 read length of metagenomic samples.

We applied HERRO (through the Dorado ONT base calling pipeline) and DeChat to the 50 Gbp human gut sample, assembled the corrected reads using each assembler, and evaluated the quality of the recovered MAGs (see Supplementary Figure S2). Assemblies incorporating HERRO consistently yielded fewer MAGs compared to raw assemblies. This likely stems from HERRO’s inability to effectively process reads shorter than 4 kbp, which are prevalent in metagenomic datasets. In contrast, DeChat significantly enhanced assembly outcomes: it improved the recovery of scMAGs by metaMDBG (11 additional scMAGs) and metaFlye (1 additional scMAG and 8 near-complete MAGs). For nanoMDBG, Dechat resulted in 2 additional scMAGs, although raw assemblies with nanoMDBG alone still produced significantly more near-complete MAGs overall (7 additional near-complete MAGs).

Although read correction can enhance assembly quality, it comes with substantial computational costs. For instance, HERRO required 56 hours (GPU mode) and 210 GB of memory, while DeChat took 30 hours and 250 GB. In contrast, nanoMDBG completed the entire assembly process, including contig polishing, in just 5 hours with 20 GB of memory. These computational costs prevented us from evaluating the read correction tools on the larger and more complex Zymo Fecal Reference and Soil data sets.

### Evaluation of metagenome assembly quality from ONT and PacBio HiFi data

#### Benchmarking setup

To enable a direct comparison between ONT and HiFi assembly quality, we generated HiFi data from the same samples used for ONT sequencing. Specifically, we produced 50 Gbp of PacBio HiFi data from the human gut sample and 250 Gbp from the soil sample, and we utilized a publicly available 255 Gbp HiFi dataset for the Zymo fecal reference sample [24] (see Table 1). We ran three HiFi metagenome assemblers: metaMDBG (v1.1), metaFlye (v2.9.3-b1797), and hifiasm-meta (v0.3-r063.2). We note that we did not apply nanoMDBG error correction to the HiFi data because HiFi reads, after homopolymer compression, a standard preprocessing step in HiFi assembly, are nearly error-free.

#### Evaluation

Figure 4a presents the number of near-complete MAGs and scMAGs recovered using both sequencing technologies at varying sequencing depths (selecting the first N reads until the desired data volume was reached). For clarity, the figure only shows results obtained with metaMDBG on HiFi data and nanoMDBG on ONT data, the combinations which consistently produced the best assemblies across experiments. Comprehensive results for all assemblers are provided in Supplementary Figure S3.

**Figure 4.**
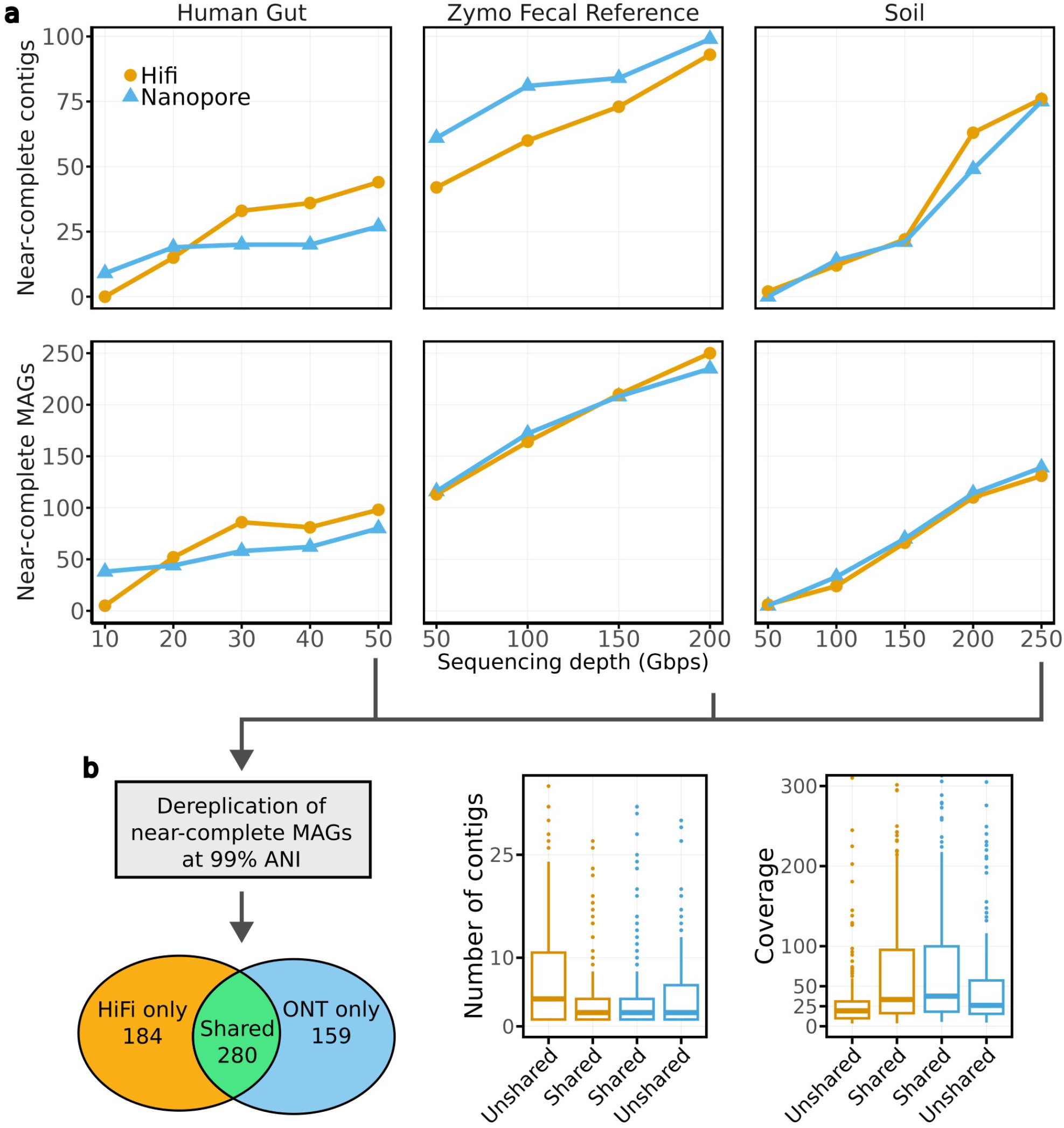
Assembly results from ONT and HiFi data generated from the same samples. For each data type, we showcase the results of the assembler which obtained consistently the best results : metaMDBG for PacBio Hifi data and nanoMDBG for ONT data. **a**, CheckM evaluation with respect to sequencing depth. We note that the number of near-complete contigs (top plots) is included in the total number of near-complete MAGs (bottom plots). **b**, Statistics of near-complete MAGs recovered by both and specific to each data type. We de-replicated MAGs per dataset using the highest sequencing depth available for each dataset.

On the human gut sample, ONT data yielded more near-complete MAGs at lower sequencing depths (33 additional MAGs at 10 Gbp of data). At this low-depth, most HiFi MAGs were of high-quality. At sequencing depths exceeding 20 Gbp, HiFi consistently recovered more near-complete MAGs compared to ONT data, with 18 more near-complete MAGs at 50 Gbp (with 20 more as single-contigs). For the Zymo fecal reference dataset, similar numbers of near-complete MAGs were reconstructed from both technologies (250 MAGs from HiFi versus 235 from ONT at 200 Gbp). However, ONT data provided more scMAGs, but with a gap that narrowed as sequencing depth increased (19 more scMAGs at 50 Gbp but only 6 more at 200 Gbp). In the soil sample dataset, the number of near-complete MAGs and scMAGs reconstructed was comparable between the technologies, with 131 near-complete MAGs from HiFi and 139 from ONT at 250 Gbp.

To evaluate differences between near-complete MAGs generated by the two technologies, we used the dRep [21] tool to identify shared and specific MAGs and assessed their coverage and contiguity (see Figure 4b). A total of 280 near-complete MAGs were shared between the technologies, while 184 and 159 MAGs were unique to HiFi and ONT, respectively. Shared MAGs generally exhibited higher coverage (median *≈* 35x versus *≈* 20x for specific MAGs) and were reconstructed in fewer contigs compared to those unique to a single technology. Unshared MAGs, therefore, represent the most challenging to assemble.

Supplementary Figure S3 explores the recovery of circular viruses and plasmids across assemblers. For the Gut sample, the number of phages and plasmids recovered was comparable between technologies (at 50 Gbp; 30 phages with HiFi using hifiasm-meta and 27 with ONT using metaFlye; 40 plasmids with HiFi using metaMDBG and 44 with ONT using nanoMDBG). In the Zymo Fecal Reference dataset, ONT consistently recovered more phages and plasmids at all sequencing depths (At 200 Gbp, 121 phages and 103 plasmids were recovered from ONT data compared to 93 phages and 85 plasmids from HiFi). For the Soil dataset, the number of recovered viruses was similar across technologies (445 with HiFi using hifiasm-meta versus 448 with ONT using nanoMDBG). The recovery of plasmids was also similar, except at the highest sequencing depth, where HiFi data recovered more plasmids (34 plasmids compared to 19). Notably, metaMDBG and nanoMDBG were the most effective assemblers for recovering circular plasmids on HiFi and ONT data, respectively. For circular phages, hifiasm-meta excelled with HiFi data, while metaFlye or nanoMDBG performed better on ONT data, depending on the sample complexity.

## Discussion

We have introduced nanoMDBG, a metagenome assembler designed for ONT long reads. Our aim was to improve upon the metaMDBG assembler by addressing the relatively higher error rates of ONT data while simultaneously preserving the high scalability, completeness and accuracy that metaMDBG achieved. We succeeded in this goal, as the read correction pre-processing step that we added in nanoMDBG only takes 6% of the total assembly run time on the large 400 Gbp soil sample. Our new correction module proved essential as we greatly improved the number of reconstructed MAGs, phages and plasmids compared to metaMDBG. Across all tested datasets, nanoMDBG outperformed metaFlye in reconstruction of near-complete MAGs, particularly single-contig ones, with the gap in assembly quality especially pronounced in complex communities. Importantly, we demonstrate that thanks to this advance, the latest ONT sequencing technology now produces results comparable to PacBio HiFi sequencing at equivalent sequencing depth, despite the remaining differences in raw read accuracy.

Given the relative cost-effectiveness of ONT reads our effective and computationally efficient assembler could prove transformative to metagenomics research. It enables the possibility of large-scale multi-sample long-read metagenomicss for simpler communities such as the human gut, which will be critical for applications such as strain tracking [26] and a fine-scale understanding of evolutionary differences over time and space. For complex communities such as soil it could be even more important, enabling for the first time large-numbers of complete genomes to be generated from single soil samples cost-effectively. However, we have also demonstrated that here results could potentially be improved. We obtained a much smaller fraction of high coverage genomes (*>*10x) that should be accessible actually as MAGs for soil, 39% vs 90% for the human gut. We are not certain of the reasons for this, it may be strain variation, but it does suggest the possibility of improved assembly strategies for these hyper-diverse communities.

In addition, to allowing the analysis of large-scale data sets, a highly efficient ONT assembler has other advantages. It makes assembly of smaller data sets feasible on for example a laptop. This aligns well with the portability of ONT sequencing, opening possibilities for on-site assembly directly in remote or field settings. In future, it may prove useful to adapt our algorithm to a streaming assembler, assembling reads in real-time as they are sequenced, the high efficiency of the underlying minimizer based denoising and assembly approach could make this feasible.

In this paper we introduce an application, nanoMDBG, that combines read error correction and assembly both in minimizer space. Our approach is far more efficient than correcting reads at the base level, i.e. sequence read error correction, and then assembling. However, we did see a slight improvement for the Gut data set in the number of single-contig MAGs when DeChat read correction was applied prior to nanoMDBG. This and the fact that the read error correction remains a resource-intensive and time-consuming task in existing long read assemblers, including state-of-the-art genomic assemblers such as hifiasm [8] and verkko [28], motivates the development of more efficient correction methods. Our minimizer-space correction approach introduces a more efficient paradigm, which we anticipate will inspire future methods to address this problem and provide fast and scalable base-level correction.

To conclude, we have shown that nanoMDBG coupled to the last ONT low error rate reads, can generate comparable numbers of high-quality MAGs as the more expesnsive but higher accuracy, HiFi PacBio reads at the same sequencing depth. This we believe will allow in the future, perhaps with additional refinements, for comprehensive genomic surveys of even the most complex microbiomes.

## Methods

Figure 1 summarises the steps of nanoMDBG, which we describe in more detail below.

### Preliminaries

We start with a lexicon of some terms and concepts related to MDBGs, assembly and sequence alignment.

#### Minimizer

In order to scale-up large datasets, our method uses only a fraction of the *k*-mers present in the reads. To select *k*-mers, we use the well-known concept of minimizers, more specifically universal minimizers [10]. Given a hash function *f* that maps *k*-mers to [0, *H*], we select a *k*-mer *m* as a minimizer if *f* (*m*) *< dH* where *d* controls the fraction (or density) of selected *k*-mers.

#### Minimizer-space read (mRead)

Prior to any treatment, reads are scanned and transformed into their ordered list of minimizers, that we call mRead. We record the positions and orientations (forward/reverse) of minimizers in each read.

#### Seed-and-chaining

The seed-and-chaining strategy is a heuristic to quickly localize potential alignments between two sequences [14, 33, 30]. Read mapping tools usually rely on this technique prior to applying a more costly base-level alignment algorithm (hence the full technique is named “seed-chaining-extend”). Seed-and-chaining is also the core procedure of nanoMDBG (but the extend step will be different).

It starts by collecting a subset of *k*-mers (the *seeds*) from the sequences to compare. In our case, we use all universal minimizers in reads as seeds. The chaining step then finds exact seed matches between the sequences (*i*.*e*. anchors) and identifies sets of colinear anchors (the *chains*). For identifying the optimal chains, we use the same scoring function and banded dynamic programming algorithm as defined in skani [33].

**Minimizer-space alignment** The seed-and-chaining procedure is usually a preliminary step prior to base-level alignment. However here, the entire alignment step will operate in minimizer-space and we will not go back to base-space. We refer to this as *minimizer-space alignment*. Sequence divergence can be directly estimated from the results of chaining as follows [5]:

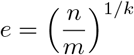

where *n* is the number of matching seeds, *m* is the number of seeds in the query and *k* is the minimizer length.

To see the difference with base-level alignment, we explain here how alignment is performed with minimizers as units (see figure 1-B) instead of bases. For a pair of mReads and matching minimizers (anchors), the other minimizers present between these anchors are examined to derive a sequence of matches, mismatches, insertions and deletions of minimizers. Such a sequence is the minimizer-space alignment. It is crucially different from standard alignment algorithms, which use minimizers but only focus on anchors and do not further try and align the other minimizers present between the anchors, they align the bases, which is more computationally expensive.

### Algorithmic challenges and motivations

The metagenomics assembler metaMDBG has been tailored to PacBio HiFi data. As input it expects long, highly accurate reads in order for its multi-k assembly strategy to be effective. In this study, we aim to extend metaMDBG to support ONT data assembly through read correction, while preserving computational efficiency to enable the assembly of large metagenomes.

Self-correction of long reads typically relies on multiple sequence alignment. This approach involves selecting each read as a target for correction, recruiting and aligning similar reads to it, and identifying correct or erroneous bases through read pileup or partial order alignment (POA) graph. This correction procedure involves an initial all-vs-all read mapping and base-level multiple sequence alignment, which are both computationally expensive. Our goal is to make this process more efficient.

Our first ingredient to speed up read comparisons is to bypass base-level alignment, performing correction at the level of minimizers only. This approach mirrors the principles of base-level correction, with mReads aligned with fast seed-and-chaining procedure instead, and minimizers stacked up to produce consensus. The corrected mReads are then assembled in minimizer-space using metaMDBG, with base-level correction deferred to the final non-redudant contigs.

Despite this efficient approach, the all-vs-all step remains computationally demanding due to the high density of minimizers required for accurate read recruitment. To address this, we introduce a second ingredient, a two-step alignment strategy tailored for read correction, based on three key observations:

1. ONT read self-correction in minimizer-space can achieve high accuracy with moderate coverage (around 20x) of recruited reads.
2. For a given target read, recruiting candidate reads that are likely to belong to same species and to the same genome location can be done quickly using the following heuristics. 1) Longer alignments are more likely to be correct than shorter alignments. 2) Alignments bounds can be estimated quickly using seed-and-chaining with low-density of minimizers.
3. Confirming that the candidate recruited reads are indeed location- and species-compatible with the target necessitates more effort as it requires to estimate sequence divergence. This step requires high-density minimizers, especially on shorter reads [33].

Our correction strategy, therefore, proceeds as follows. For each target read, we recruit the most similar reads (up to approximately 20x coverage) using seed-and-chaining with a very-low density of minimizers. We then remove divergent reads using a higher density of minimizers. This strategy restricts the more costly divergence analysis to a subset of reads, all read vs approximately 20x coverage, rather than requiring it for all reads. The following section describe our correction strategy and heuristics in more details.

### Read correction in minimizer-space

The following steps describe the novel correction step specifically (see figure 1-C). In a nutshell, we first perform a fast all-vs-all read mapping using a low density of minimizers (0.5% by default). The output of this step is, for each read, the list of their best read matches (up to 20x reads). For a given target read to correct, we recompute its alignments using a higher density of minimizers (2.5% by default), and filtered recruited reads more accurately. We then pileup the minimizers using a variation graph and determine the most supported path as consensus. We describe each of these steps more precisely in the following.

#### Converting to minimizer-space

We start by converting the reads into their ordered list of universal minimizers, that we call mReads. We extract two sets of mReads. One with a low-density of minimizers (0.5% by default) which will be used to quickly perform all-vs-all read mapping. And another with a high-density of minimizers (2.5% by default) which will be used to accurately correct mReads.

#### Ultra-fast mRead recruitement using low-density minimizer-space alignment

In this step, we perform all-vs-all read seed-and-chaining in order to find for each read, its best matching reads that will be used for correction. In order to speed-up this costly process, we use a low density of minimizers (0.5% by default). We use the following simple formula to rank alignments : score = number of minimizer matches - number of differences, where the number of differences is the sum of missmatching miminizers, deletions and insertions. For each read, we recruit up to 20x reads. Other alignments are discarded.

#### Accurate high-density alignment filtering

In this step, we perform seed-and-chaining alignment between the target mRead and the recruited mReads using their full minimizer representation (2.5% by default). Alignments with more than 4% divergence are discarded in order to discard reads that belong to other species while allowing for a certain error rate. Additionally, we discard alignments smaller than 1000 bps or with over hang longer than 2000 bps, in order to correctly recruit reads in genomic repeats.

#### Piling up minimizer with minimizer-space variation graph from high-density alignments

After accurate alignment filtering, we start the construction of a variation graph from the remaining alignment (see figure C-3). We initiate the graph with the minimizers of the target mRead. We then process sequentially each high-density alignment computed in the previous step. A node is added to the graph when a mismatch or an insertion occur (see respectively node “m7” and node “m8” in figure 3-C). We also record the support of each minimizer transition.

We note that this is a simpler approach than a partial order graph (POA) since the alignments are only perform against the target mRead, instead of the against graph in the case of the POA. We found that the correction with this strategy was good enough for assembly. It has the advantages of being faster than the POA and the resulting variation graph does not depend on the order in which sequences are added to the graph.

If a FASTQ file is provided as input, we weight edges by quality scores instead of raw transition counts. The quality score of a minimizer is defined as the minimum quality score among its constituent bases. Each edge in the graph is then weighted by the sum of qualities of its source and destination minimizers.

#### Generating consensus

After all alignment has been added to the variation graph. We extract the most supported path from the graph as consensus.

### Assembling corrected minimizer-space reads

The corrected mReads are assembled using metaMDBG [3], with only minor modifications for the conversion of contigs from minimizer-space to base-space. A brief summary of the methods of MetaMDBG is provided here for completeness. MetaMDBG uses the minimizer de-Bruijn graph as a core structure. It assembles mReads using an efficient multi-k strategy in minimizer-space for handling uneven species coverage. At each multi-k iteration, an abundance-based algorithm termed ‘local progressive abundance filter’ is applied to remove potential inter-genomic repeats, strain variability and complex error patterns. The resulting minimizer-space contigs (mContigs) are finally converted into their base-space representation. Previously, our method for reconstructing mContig sequences was designed for highly accurate reads. However, for ONT data, we developed a more adaptable approach. We map the raw reads (in minimizer representation) to the mContigs using the seed-and-chaining algorithm that we developed for our correction module. We then identify the longest alignments and extract the sequences from the raw reads that are spanned by the minimizers to reconstruct the mContig sequences. The final draft assembly is polished using a racon-like [37] approach.

### Metagenome sequencing data generation

Detailed protocols for DNA extraction and platform specific library preparations are available at (https://www.protocols.io/view/soil-metagenome-pacbio-djga4jse.html and https://www.protocols.io/view/soilmetagenome-ont-dhrm3546.html). A summary of sample collection, DNA extraction and library preparations are provided here.

#### Sample collection and storage

All samples were collected and stored in Zymo DNA/RNA shield and left to incubate at 4 oC for three hours prior to snap freezing and storage at -80 oC prior to DNA extraction.

Soil was collected using a sterile soil corer and passed through a sterile soil sieve to homogenise the sample. A total of 10g of homogenised soil sample was mixed with 100 ml of Zymo DNA/RNA shield and distributed into 1 ml aliquots containing 100mg/ml of soil.

Faecal samples were provided as part of an ongoing human study “QIB colon model study”. The study was approved by the local Quadram Institute Bioscience Human Research Governance committee (IFR01/2015) and by the London-Westminster Research Ethics Committee (15/LO/2169). The trial was registered at clinicaltrials.gov (NCT02653001). All participants provided signed informed consent prior to donating samples. The study was conducted in accordance with the Declaration of Helsinki.

Faecal samples are provided by volunteers of both genders, aged between 25 and 54. All participants declare to be in good health, have no diagnosed chronic gastrointestinal health problems, and have not consumed antibiotics for at least six months prior to stool donation. Donors are not instructed to follow any diets or consume any specific foods, in accordance with the terms of the ethical approval.

A total of 10g of fresh faecal material was processed in a stomacher with a total of 100 ml of anaerobic PBS. Faecal slurry was then collected and distributed into 1 ml aliquots. Aliquots were centrifuged for 10 min at 10,000 x g at Room temperature and the supernatant discarded. Samples were then homogenised with 1 ml of Zymo DNA/RNA shield and stored at -80 oC prior to DNA extraction.

#### DNA Extraction

DNA was extracted from all samples using the MPBio Fast DNA spin kit for soil with slight modifications from the manufacturer protocol. Modifications include processing samples on ice, a reduced homogenisation time of 2 × 10 sec at 5.0 m/s with 5 min of cooling between runs. DNA was eluted first into 100 ml of DES elution buffer at 56 oC with the addition of 1 ml of RNaseA. Samples then undertook at 0.5 X SPRI clean up prior to both PacBio and ONT library preparations.

#### PacBio Library preparation and sequencing

PacBio libraries were constructed using SPK 2.0 kit and the overhang SMRT bell adapter. Samples were size selected using a 3.7 X dilute SPRI cleanup prior to sequencing. Libraries were sequenced using the PacBio Revio device and high-density flow cells and ran for a total of 24 hour movie time with a hifi q score cut off at q20.

#### Nanopore Library preparation and sequencing

Nanopore libraries were constructed using LSK-114 and processed using the long fragment buffer workflow. Libraries were sequenced on a P2 solo device using R 10.4.1 flow cells (Flo-Pro 114M). Samples were sequenced for a total of 96 hours with a score cut off at q10. Sequences were base called and trimmed using Dorado SUP V5 basecalling model in post processing.

### Assembling data sets, mapping reads and binning contigs

We ran all assemblers with 32 central processing unit threads. We ran metaMDBG with option ‘–in-ont’ for ONT data (to activate nanoMDBG method) and with the option ‘–in-hifi’ for HiFi. We ran metaFlye with the options ‘–meta’ and ‘–nano-hq’ for Nanopore data sets and with option ‘–pacbio-hifi’ for HiFi data sets. We ran hifiasm-meta with default parameters. We only used the hifiasm-meta primary assembly of polished contigs (p ctg.gfa), as adding alternate contigs reduced the overall MAG quality. We used the command ‘/usr/bin/time -v’ to obtain wall-clock runtime and peak memory usage. All tools that were used and the complete command line instructions are available in Suplementary Table 1.

We mapped reads to contigs using ‘minimap2’ with option ‘-x map-ont’ for ONT reads and with option ‘-map-hifi’ for HiFi reads. We filtered out reads in which all of the alignments were shorter than 80% of its length, and we assigned each remaining read to a unique contig through its longest alignment (breaking ties arbitrarily). We performed contig binning using SemiBin2 [22] (v.2.1.0), using ‘single easy bin’ command with options ‘–sequencing-type=long read’, ‘–self-supervised’ and a fixed seed (–random-seed 42) for reproducibility.

### Quality assessment of assemblies

We used CheckM2 (v.1.0.1) to assess the quality of all MAGs. We used Barrnap (https://github.com/tseemann/barrnap), and Infernal [19] to predict, respectively, rRNA and tRNA genes from near-complete MAGs. We filtered out annotations with E-values over 0.01. We denoted near-complete MAG as RNA complete if they have at least one full-length copy for all three types of rRNAs, and at least 18 full-length copies of tRNAs. We used genomad (v.1.8.0) using command ‘end-to-end’ with options ‘–conservative’ to identify strongly supported plasmids and viruses in each assembly. We used checkV [20] to assess the quality of viral contigs.

### Assessment of completeness and fragmentation of assemblies with reference sequences

We used the following process to assess the completeness and fragmentation of assemblies when reference genomes are available (mock reference genomes). We used wfmash to align contigs against the reference sequences. Alignments with less than 99% identity were filtered out. Alignments were ordered by their matching score *MS* = *alignLength alignIdentity* (best score first). We considered alignment identity to improve contig assignment to similar strains. Alignments were then processed sequentially and contigs were uniquely assigned to references. During this process, we check whether a reference is complete or not, meaning that at least 99% of its positions are covered by contigs. We prevent other contigs from being assigned to a complete reference. Moreover, we prevent a contig to be assigned to a reference if more than 30% of its matching positions are already covered by another contig. In this case, we first try to assign this contig to another reference. References with less than 70% completeness were considered missed by the assembler.

### Taxonomic classification of MAGs recovered from the ONT soil sample

The phylogenetic tree of Figure 2c was built using fasttree [25] from the output alignment of gtdbtk version 2.1.0 [7] on near-complete quality MAGs of all three assemblers for the anaerobic digester dataset. Concurrent diversity coverage between the different assemblers was explored at different taxonomic levels from genus to domain. To do so, it is necessary to first address MAGs for which no annotation is available at a given taxonomic rank. A pair of unannotated MAGs may or may not share the same taxa. A first pass based on tree topology allows us to select neighbouring MAGs as candidates for sharing the same unknown taxa. As a second step we compute the Relative Evolutionary Distance using the R library Castor version 1.7.3 [16]. Following guidelines from gtdb, we use their median RED values for each taxa in order to decide on grouping together unknown MAGs. We then find the best ancestor for each unknown MAG in terms of their RED being nearest to the corresponding taxa median RED. If they share the same best ancestor, we group them together otherwise we split them into distinct unknown taxa. Tree manipulation and representation is carried out using the library ggtree version 2.4.1 [41], treeio version 1.14.3 [38] and ggtreeExtra version 1.0.2 [40].

### Identification of Single-Copy Core Genes (SCGs)

In order to assess the species diversity of the assemblies and hence quantify our ability to capture this diversity as MAGs, single copy core genes (SCGs) were identified from contigs. First, open reading frames (ORFs) were called using prodigal [12] (v2.6.3) with option -p meta. A set of 36 different SCGs were annotated using RPS-BLAST (version 2.16.0) [2] using the pssm formatted COG database [36], which is made available by the CDD [17]. Annotation was performed with a best hit strategy, filtering out any hit with an e-value higher than 1e-10 and for which the alignment represents less than 50% of the subject length. Resulting hits were de-replicated at 97% using vsearch (2.29.1) [29] with option -cluster smallmem. Species diversity was estimated as the median abundance across all SCGs.

A custom Python (version 3.10) script was used to link previously computed contig coverages, inclusion of contig in a MAG, related MAG quality and SCGs. Coverage of SCGs from the same cluster were summed and when multiple MAG qualities could be found, the higher one was taken. This resulted in a table stating for each SCG cluster, its coverage and if it was found included in a MAG as well as related quality. This table was plotted using the ggplot2 [39] package from R (version 4.3.3) to give Figure 3.

## Data availability

The sequencing read data generated for this study are available at ENA bio project PRJEB88618; accession numbers are given in Table 1. Zymo mock reference genomes are available at https://s3.amazonaws.com/zymo-files/BioPool/D6331.refseq.zip. The ONT Zymo Fecal Reference data set is available https://epi2me.nanoporetech.com/lc2024-datasets/. The HiFi Zymo Fecal Reference data set is available at https://www.pacb.com/connect/datasets/#metagenomics-datasets.

## Code availability

We implemented the nanoMDBG method in the metaMDBG software (https://github.com/GaetanBenoitDev/metaMDBG). The nanopore mode is activated using the input parameter (–in-nano), and the original PacBio HiFi mode using parameter (–in-hifi). The analysis scripts used in this study to compare assemblers are available at https://github.com/GaetanBenoitDev/NanoMDBG_Manuscript.

## Acknowledgements

C.Q. and S.R. acknowledge the support of the Biotechnology and Biological Sciences Research Council (BBSRC), part of UK Research and Innovation; Earlham Institute Strategic Programme Grant (Decoding Biodiversity) BBX011089/1 and its constituent work package BBS/E/ER/230002C; the Core Strategic Programme Grant (Genomes to Food Security) BB/CSP1720/1 and its constituent work packages BBS/E/T/000PR9818 and BBS/E/T/000PR9817; and the Core Capability Grant BB/CCG2220/1. We acknowledge the assistance of Dr. Susheel Bhanu Busi (CEH, Wallingford) in helping to organise the soil sampling.

## Supplementary Figures

**Figure S1.**
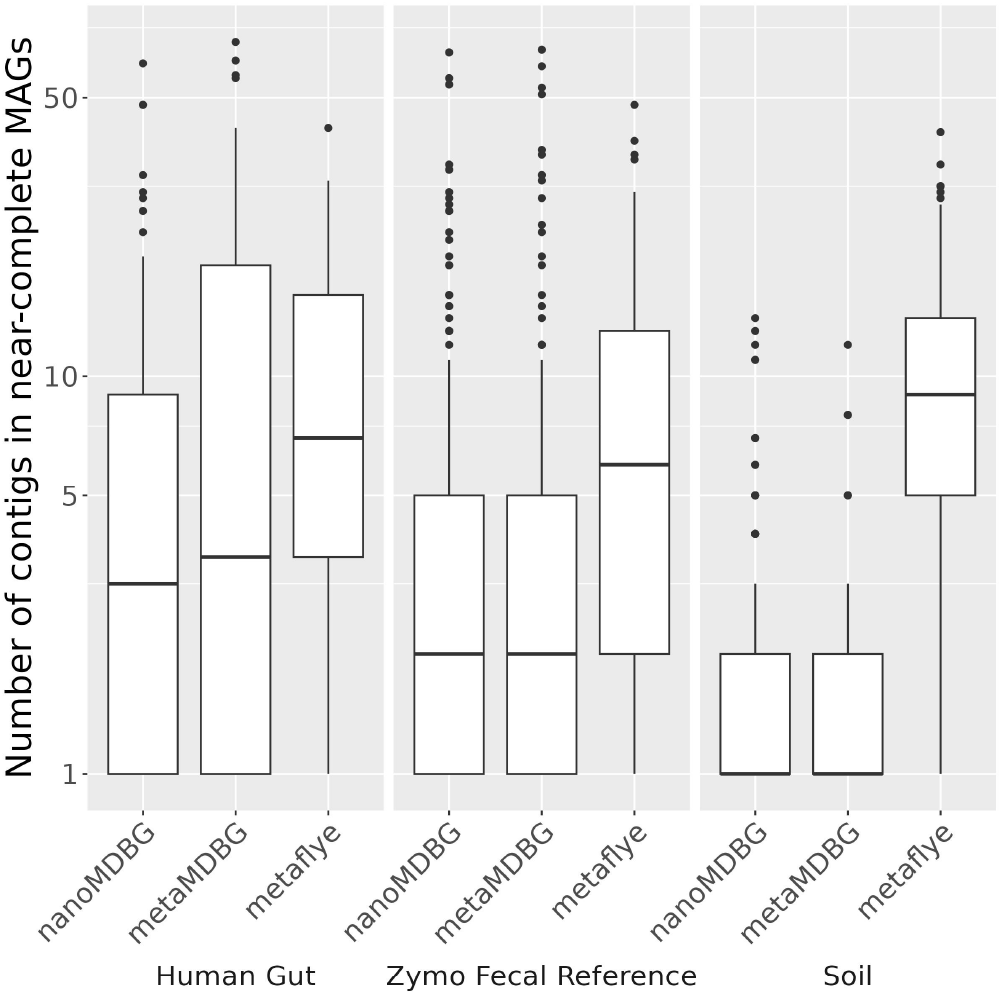
Number of contigs in near-complete MAGs across the three ONT samples. The boxplot elements are the median (horizontal bar), 25th and 75th percentiles (box limits Q1 and Q3), Q1-1.5*IQR and Q3+1.5*IQR (whiskers, IQR=Q3-Q1) and outliers. Summary statistics (n, min, median, mean, max): Human gut - nanoMDBG (80, 1, 3, 7.8, 61); metaMDBG (50, 1, 3.5, 12.8, 69); metaFlye (59, 1, 7, 10.6, 42) : Zymo Fecal Reference - nanoMDBG (235, 1, 2, 5.3, 65); metaMDBG (179, 1, 2, 5.6, 66); metaFlye (167, 1, 6, 8.8, 48) : Soil - nanoMDBG (296, 1, 1, 2, 13); metaMDBG (52, 1, 1, 2, 12); metaFlye (116, 1, 9, 10.5, 41).

**Figure S2.**
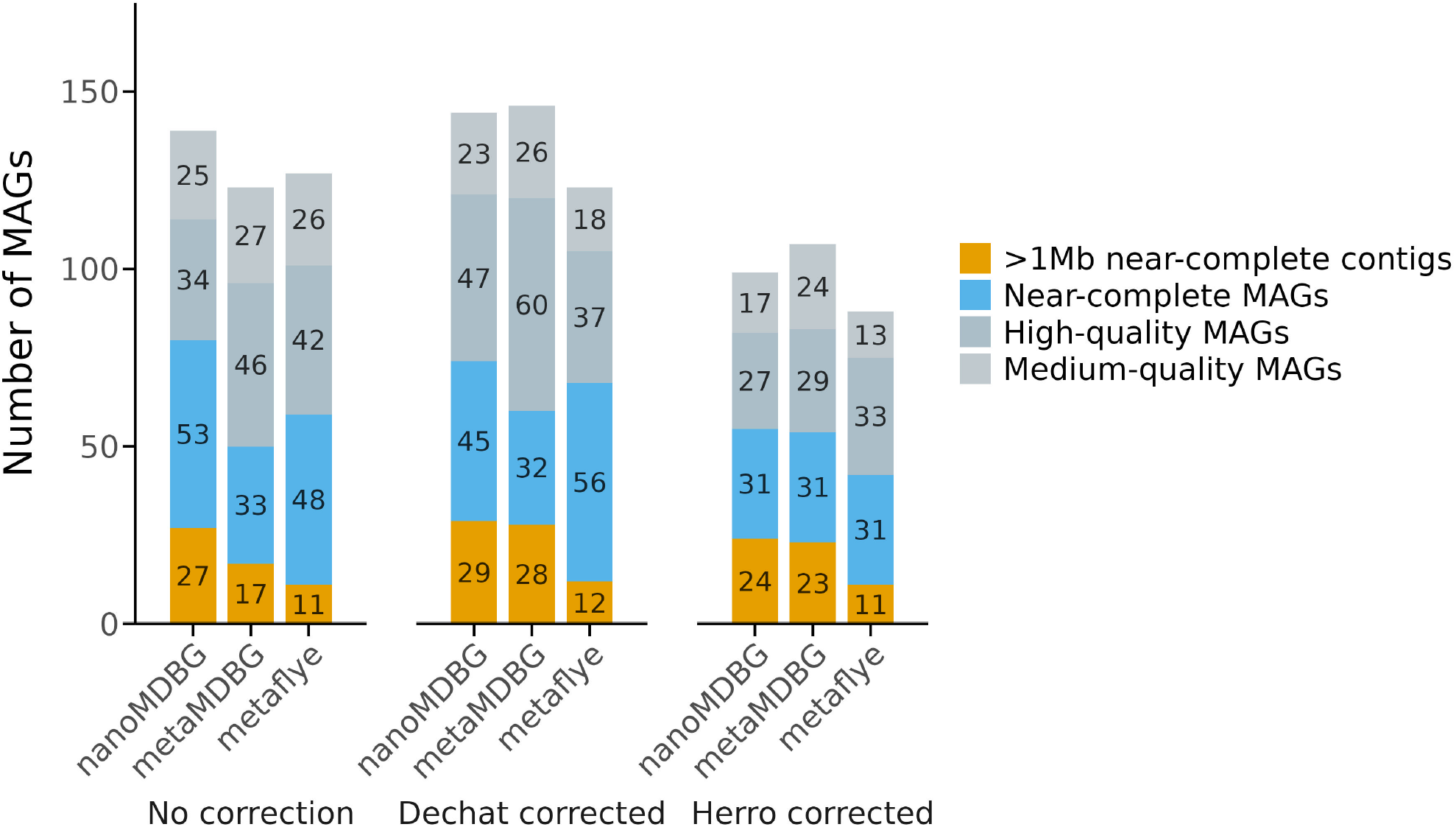
Assembly results on ONT Human gut sample using preliminary base-level correction methods. We ran dechat and herro correction tools on the 50 Gb Human gut sample, followed by assembly with nanoMDBG, metaMDBG and metaFlye. “No correction” corresponds to the assembly results on the raw read sets.

**Figure S3.**
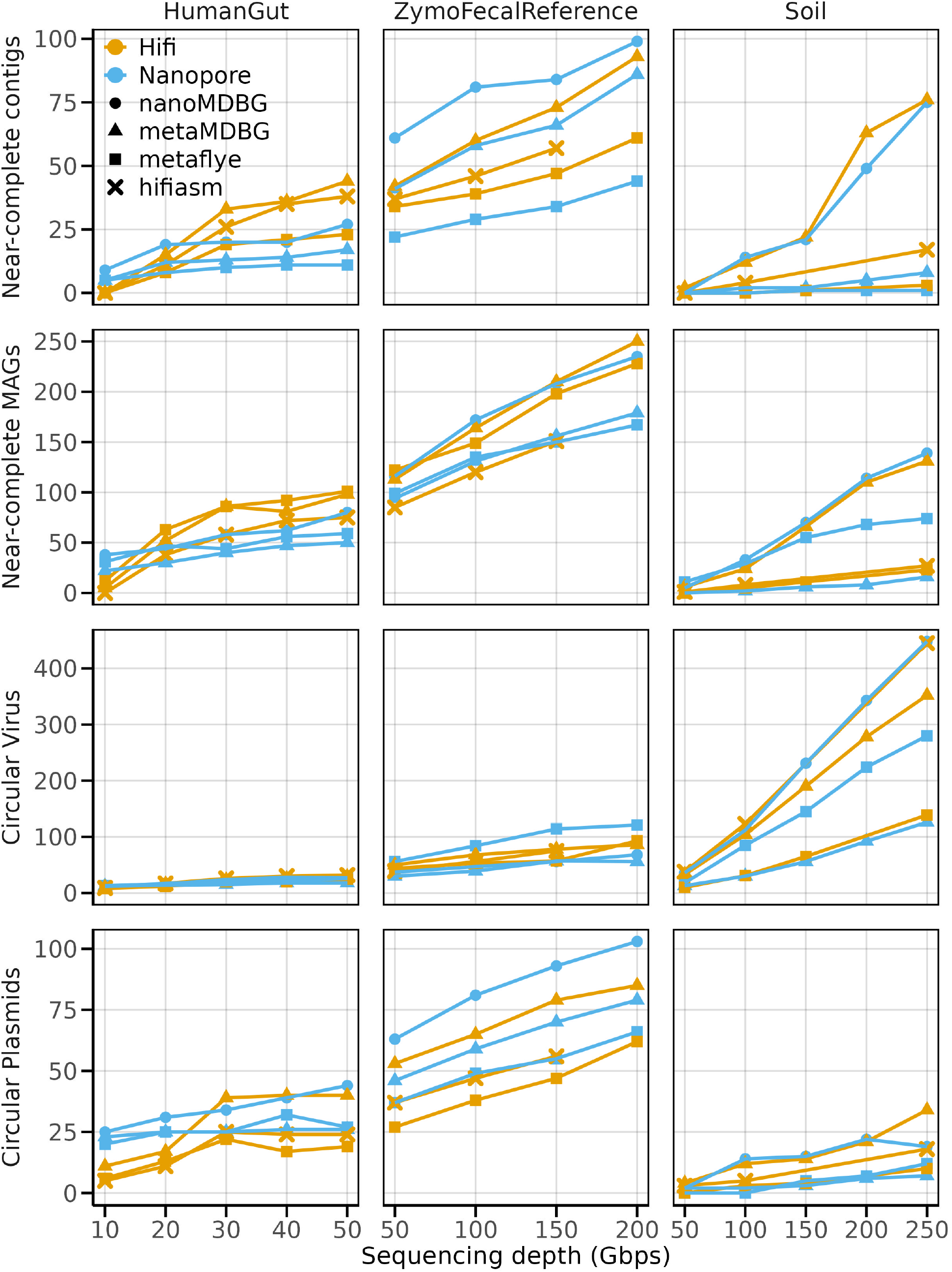
Assembly results for every assemblers from ONT and HiFi data generated from the same samples

